# Biogeographic history and megafauna shape frugivory interaction networks across equatorial forests

**DOI:** 10.64898/2026.08.12.744530

**Authors:** G Clément, N Lotfi, L Ong, A Campos-Arceiz, F Bretagnolle, KR McConkey, R Thomachot, MAR Mello, P-M Forget

## Abstract

Frugivory is essential for the maintenance of tropical forests, influencing seed movement, spatial structure, and ecosystem functioning. Although frugivory interactions are well documented at local scales, we still know relatively little about how network structure varies across biogeographic regions, particularly in equatorial lowland forests. In this study, we compare frugivory networks in lowland forests in the Guiana Shield (South America), Malaysia (Southeast Asia), and Central Africa (Gabon and Cameroon). Because these sites occur at similar latitudes, they provide a useful setting to examine how evolutionary history and biogeographic context, rather than climate alone, might shape the structure of the network. We analyze patterns across regions, focusing on implications for frugivory interactions and, ultimately, seed dispersal. Guided by the Integrative Hypothesis of Specialization (IHS), we focus on the role of megafauna and other large-bodied frugivores, which are important consumers of large fruits and dispersers of large seeds, and have experienced different historical trajectories in each continent. We also considered whether smaller frugivores, such as rodents, might partially compensate for the loss of Megafauna in the Americas through seed handling and caching behavior. The pantropical comparison revealed structural and functional differences between the studied frugivory networks. In the Guianas, lacking megafauna, the frugivory network showed strong modularity with nestedness within modules, forming a marked compound topology. Alternatively, in Central Africa, with elephants and great apes, the network showed intermediate modularity and nestedness, reflecting a balance between local compartmentalization and regional integration. Finally, in Malaysia, the network was predominantly nested, with a few highly connected species, especially figs and flying foxes, that linked most of the partners. Across continents, frugivory networks tend toward a compound topology, but the relative influence of modular and nested components shifts with biogeography, evolution, and ecology.

## 1 Introduction

Ecological interactions are the glue that binds different plant, animal, and microbe species, sustaining biodiversity and ecosystem services (Stanworth et al., 2024). Among these interactions, frugivory and seed dispersal play a central role in maintaining ecosystem functioning, as they influence regeneration, stability, and adaptation to environmental change (Dennis et al., 2007). In tropical forests, where plant species coexist in high densities, dispersal is critical for plant recruitment (Clark and Clark, 1984). By moving seeds in space, frugivores ultimately connect plant populations and shape the spatial and genetic structure of entire communities (Levine and Murrell, 2003). In addition, being a key to community assembly, frugivory is also diverse and widespread: between 75 and 90% of all tropical plant species, from trees to lianas, produce fleshy fruits that animals consume (Landim et al., 2024). These interactions involve ants, bats, birds, carnivores, fishes, lizards, primates, rodents, snails, and many other animals that disperse seeds in a variety of ways (Forget et al., 2007; McConkey et al., 2024). Large-bodied frugivores disperse the widest range of seed sizes, including very large seeds (Forget et al., 2007). Primates often discard seeds after consuming their pulp (Sivault et al., 2023). Bats defecate or drop small seeds over large distances and can also carry heavier seeds and fruits (Lobova et al., 2009), (Villalobos-Chaves et al., 2020). Many plant and animal traits influence their interaction success, as well as potential seed-dispersal outcomes (Sivault et al., 2020), (Ong et al., 2022). Phenology also influences interaction probabilities, as an animal cannot consume a plant that bears fruit outside its activity season. Similarly, species whose ranges do not overlap cannot interact with each other. Consequently, these morphological, temporal, and spatial constraints give rise to “forbidden links” that restrict the interactions within a network (Duchenne et al., 2025). Additionally, the taxonomic diversity of plant and frugivore communities, which results in particular from the evolutionary and ecological history of their environment and from stochasticity, also influences who is linked to whom (Carlucci et al., 2017).

These patterns in frugivory and potential seed dispersal scale-up from individuals to entire communities, and are observed across different spatiotemporal scales (Levine and Murrell, 2003). Despite this multiplicity, large-scale frugivory patterns remain poorly understood (Mello et al., 2019). Regional differences in evolutionary and climatic histories shape plant and frugivore assemblages, which, in turn, shape how they are interlinked (Moulatlet et al., 2023). Tropical America, for example, shows greater diversity of fruiting plants and small frugivores than Africa, likely a function of plant dispersal by floatation across the Atlantic ocean Bardon et al. (2013, 2016); Dick and Heuertz (2008) and higher angiosperm diversification between continents, although Africa still retains a richer assemblage of large-bodied frugivores (Carlucci et al., 2017). Regarding latitudinal gradients of diversity, some studies point to stronger network specialization in American forests (Dugger et al., 2019), whereas others report a decrease in specialization in the tropics (Schleuning et al., 2012). Latitudinal gradients are highly complex and controversial in other interactions, such as pollination Sakhalkar et al. (2025); Moles and Ollerton (2016). Anyway, equatorial lowland forests are still underrepresented and Southeast Asia has rarely been directly compared to Africa or America, when it comes to frugivory studies. Tropical forests in South East Asia have been largely underrepresented in comparative studies of plant–frugivore networks because relatively few networks in this region have been made available. As a result, the structure of Southeast Asian networks has still not been rigorously compared with those of Africa and the Neotropics (Schleuning et al., 2012).

In addition to addressing this Wallacean shortfall Hortal et al. (2015), integration of network science into large-scale studies on species interactions is one of the key challenges today, as the idea that they may exhibit a biogeographical structure is often overlooked (Windsor et al., 2023). Furthermore, patterns of diversity in different tropical biogeographic regions are influenced by their evolutionary and climatic history, which leads to variation in the taxonomic composition of their plants and animals. Such historical differences might influence the properties of frugivory networks within these different biogeographical regions. Thus, a higher degree of specialization among frugivores was demonstrated within North American networks (Dugger et al., 2019). On a more global scale, evidence points to latitudinal variation in the structure of mutualistic plant-frugivore interaction networks (hereafter, frugivory networks), with a decrease in specialization towards tropical latitudes, a pattern also observable across Africa and the Americas (Schleuning et al., 2012). Despite these intriguing initial findings, which highlight the importance of studying the biogeography of interaction networks, finer gaps remain. In particular, lowland tropical forests are absent from most studies (Schleuning et al., 2012). Furthermore, previous studies focused primarily on East African forests, which are subject to a drier climate, so the frugivory networks sampled were predominantly from savannas, which may have introduced biases when comparing them with networks from South American forests (Dugger et al., 2019). Finally, frugivory networks in the tropical forests of Southeast Asia are underrepresented and their structure has not yet been compared with that of African and American networks, especially from a historical perspective (Schleuning et al., 2014).

To address these gaps, we worked under the framework of the Integrative Hypothesis of Specialization (hereafter IHS) (Mello and Dormann, 2025; Mello et al., 2025), and tested whether predictions deduced from this hypothesis hold for frugivory networks on a biogeographic scale. Such a large scale has rarely been investigated in ecological network studies (Nakamura et al., 2024), but interest is growing, as it is more suitable for assessing evolutionary processes and fundamental niches (Fründ et al., 2016). Here, we compare frugivory networks from equatorial lowland forests in the Guianas (South America), Peninsular Malaysia (Southeast Asia), and Gabon and Cameroon (Central Africa). These sites share similar latitudes, minimizing biases from potential latitudinal gradients in network structure (Schleuning et al., 2012). We hypothesized that American networks should exhibit a higher degree of specialization and stronger niche partitioning among frugivores than their African and Asian counterparts. This pattern should result from the historical loss of megafauna dispersers in the Neotropics, including gomphotheres and other large-bodied mammals that disappeared during the Late Pleistocene (Guimarães Jr et al., 2008). In contrast, tropical forests in Central Africa and Southeast Asia still retain large frugivores such as elephants and great apes, which are known to play a central role in the dispersal of large seeds and the maintenance of network connectivity (Albert-Daviaud et al., 2022), so they are expected to display more cohesive and less modular networks. In addition, we examined whether smaller frugivores, particularly rodents, might partially compensate for the ecological functions formerly performed by megafauna through synzoochory and seed-caching behaviors, as such compensation mechanisms may be especially important in regions where large vertebrate dispersers are absent. This expectation is supported by previous work that highlights the importance of fruit and seed handling capacity in shaping frugivory networks (Ong et al., 2022). Through this comparative approach, we aim to determine whether frugivory network structure differs consistently among the three major tropical regions and to identify the ecological and evolutionary processes underlying these patterns. In particular, we assess whether historical megafauna loss in the Neotropics and differences in trait matching between plants and frugivores have contributed to continental-scale variation in network organization. Finally, we test whether the Integrative Hypothesis of Specialization provides a useful framework for explaining the topology and species roles of tropical plant–frugivore networks across continents.

## 2 Methods

### 2.1 Data

Three datasets were provided to us by various collaborators, from which we were able to generate three equatorial networks in order to carry out a transcontinental comparative study: a network based on data from the Guianas (South America), a network combining data from Gabon and Cameroon (Central Africa), and a third network located in Malaysia (Southeast Asia), Table. 1. These networks represent frugivory interactions recorded empirically in the field and using local ecological knowledge from indigenous people, including fruit consumption and fruit or seed handling events, rather than direct measures of seed-dispersal effectiveness. Therefore, throughout the manuscript we interpret network structure as the architecture of frugivory interactions, while discussing seed-dispersal consequences only as potential ecological implications.

**Table 1:** Summary of study regions, study sites, interaction numbers, plant species, and animal species used in this study.

| Region | Country | Main Study Sites | No. of Interactions | No. of Plant Species | No. of Animal Species | Reference |
| --- | --- | --- | --- | --- | --- | --- |
| America | Guiana Plateau | Cabassou Forest, Nouragues Nature Reserve, Paracou (CIRAD), Saint-Elie Trail, Saint-Eugène Scientific Station, Brownsberg Nature Park, Lake Guri, Raleighvallen-Voltzberg Reserve | 1807 | 660 | 62 | (Forget et al., 2007) |
| Africa | Complete database | All African records before filtering | 12204 | 844 | 347 | (Durand-Bessart et al., 2023) |
| Africa | Gabon | Makokou, La Lopé, Belinga, Loango, Lékédi, Moukalaba-Doudou | 2700 | 325 | 96 | (Durand-Bessart et al., 2023) |
| Africa | Cameroon | Dja Forest | 715 | 205 | 19 | (Durand-Bessart et al., 2023) |
| Asia | Malaysia | Royal Belum State Park | 2052 | 162 | 34 | (Ong et al., 2022) |

An initial screening was carried out for the African database. Study sites with a substantial number of observations were selected to avoid possible biases related to the sampling effort. In Gabon, the Makokou and Lopé sites account for almost all the interactions observed in that country (Durand-Bessart et al., 2023). However, other sites with a small number of observations were retained, as they contain information on interactions involving several specific groups of primates rarely observed elsewhere. These include gorillas and chimpanzees in Belinga and Loango, mandrills in Ĺekédi, and various other groups in Moukalaba-Doudou. In Cameroon, only data from the Dja Forest were used, as they record 715 interactions compared to fewer than 100 at all other stations. Due to their proximity and to complete the African ‘equatorial network’, all data from these seven sites were merged into a single network named ‘Central Africa’, Fig. 1.

**Figure 1:**
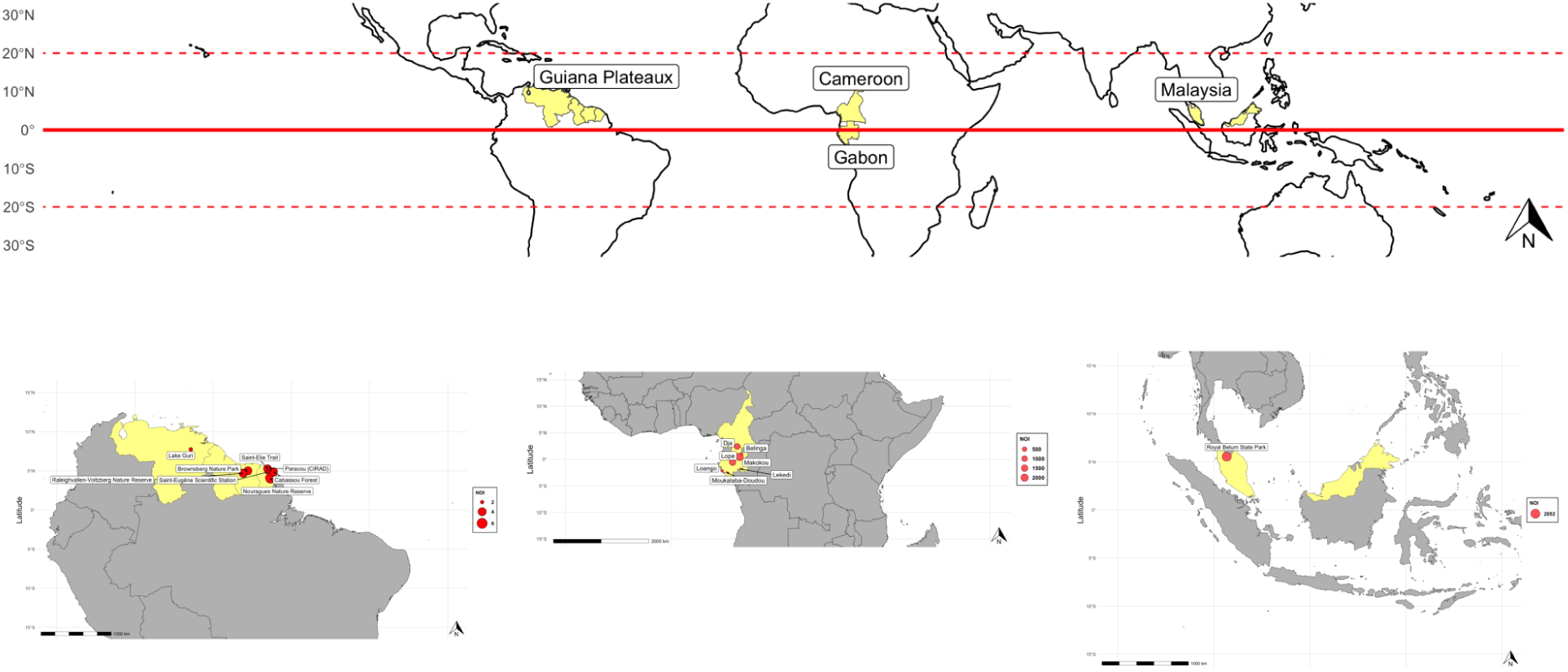
Locations of study sites where frugivory interactions were sampled to construct the three equatorial networks. From left to right: Guianas, Central Africa (Cameroon-Gabon), and Malaysia

Because the three databases were compiled from different sources, they also differ in sampling history, taxonomic resolution, and detectability of frugivore groups. The Malaysia network, in particular, is not fully resolved at the species level for all interacting taxa, as several animals are represented by groups of closely related species, such as rats, squirrels, hornbills, bats, and bulbuls (Ong et al., 2022). Rodents and nocturnal vertebrates are difficult to identify consistently across datasets, and some groups are identified at the family or functional-group level. These differences are addressed by using binary matrices and identical network metrics for all regions, but they must be considered when interpreting cross-regional differences in modularity, nestedness, and centrality.

### 2.2 Topological analysis

All network and statistical analyses were carried out using R version 4.6.0. The data were first transformed and then analyzed using the *tidyverse* (Wickham et al., 2019), *igraph* Csárdi and Nepusz (2006), and *bipartite* packages (Dormann et al., 2009), in addition to user-defined functions (see Supplementary Materials).

Interaction data were converted into adjacency matrices. To ensure consistency, the matrices followed the same structure in all datasets: columns represented species at the higher trophic level (frugivores), rows represented taxa at the lower trophic level (plants), and entries were binary (0 = absence of interaction, 1 = presence of interaction). The binary format was adopted because the data were compiled from heterogeneous sources using different sampling methods, often lacking standardized information on interaction frequency or species abundances (a very common limitation, see (Kita et al., 2022)).

The resulting networks had their topology examined under the framework of IHS (Mello et al., 2025; Mello and Dormann, 2025), which proposes that a balance between interaction constraints and neutral processes shapes topology. Here, we consider the four archetypal topologies hypothesized in a seminal paper: nested, modular, gradient, and compound (Lewinsohn et al., 2006). It is important to note that frugivory networks are commonly regarded as mutualistic, although, in fact, they contain not only positive interactions, but also negative (Carrete et al., 2022; Simmons et al., 2018) and dual interactions (Genrich et al., 2017). It has been generally assumed that most mutualistic networks would have a nested topology, in other words, the interactions of the least connected species would represent a subset of the interactions made by the most connected species (Bascompte et al., 2003). Nevertheless, growing evidence indicates that mutualisms also form modular networks, in which species are organized into cohesive subgroups of densely connected partners (Fortuna et al., 2010). In addition, recent evidence indicates that, when properly sampled, taxonomically comprehensive, local mutualistic networks may present a compound topology, which combines a modular structure with internally nested modules (Lampo et al., 2024). Gradient topologies, in other words, networks with only a few realized links and very little niche overlap, have been seldom detected or even investigated in nature (Mello et al., 2025).

Given this complex scenario of archetypal topologies, we conducted a thorough structural assessment using a protocol that combines multiple network metrics with null model analysis (Felix et al., 2022; Queiroz et al., 2021), as detailed in the following sections.

### 2.3 Modularity

Modularity quantifies the degree to which a network is divided into cohesive subgroups (i.e., modules), where the density of connections is greater within the same module than between different modules (Newman, 2004). It also provides a way to identify modules in which link density is higher among nodes of the same module than between nodes of different modules (Olesen et al., 2007). We quantified modularity using the Q metric and identified modules with the computeModules function of the bipartite package, based on the DIRTLPAwb+ algorithm (binary version) (Beckett, 2016).

### 2.4 Nestedness

Nestedness measures the extent to which the interactions of nodes with lower degrees represent subsets of the interactions of nodes with higher degrees (Ulrich et al., 2009). We used the NODF metric to quantify binary nestedness (Almeida-Neto et al., 2008). Because modular networks confound nestedness across network structural levels (i.e., entire network, layers, and modules), nestedness was partitioned into two components: nestedness among nodes of the same module (NODF*_SM_*) and nestedness among nodes of different modules (NODF*_DM_*) (following (Felix et al., 2022)).

### 2.5 Null model analysis

To test for the significance of observed modularity (Q) and nestedness scores (NODF, NODF*_SM_*, NODF*_DM_*), we generated 1,000 randomized matrices using the free null model (equiprobable) and the restricted null model (degreeprob) implemented in the package *bipartite*. Values significantly higher or lower than the corresponding null distributions indicate that the observed network structure deviates from expectations in random assembly, with the restricted model additionally accounting for the degree distributions of species. Statistical significance was evaluated using Student t-tests.

### 2.6 Z-scores

For cross-network comparisons, observed values were standardized as Z-scores (Eq. 1) by subtracting from them the mean and dividing the difference by the standard deviation of the values measured for randomized networks produced with null models.

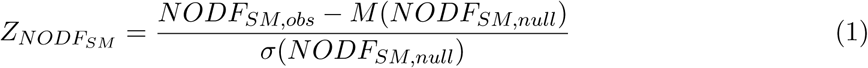

### 2.7 Species-level analysis

To characterize the functional role of frugivore species within modular networks, we calculated within-module (*z*) and between-module connectivity (*c*) (following (Olesen et al., 2007)). Based on threshold values (*z* = 2.5, *c* = 0.62), species were classified in the following functional roles: peripherals, module hubs, connectors, or network hubs.

## 3 Results

### 3.1 Guianas

The Guianas network was the most species-rich among the three, comprising 660 plant species, 62 frugivore species, and a total of 722 taxa. Both assemblages were compositionally unique, as indicated by the high Whitaker beta-diversity relative to the other regions (*β_plant_*= 0.99 with Central Africa; *β_plant_* = 1.00 with Malaysia; *β_frugivore_* = 1.00 with both). These values confirm the strong taxonomic turnover of the Guianan frugivory system, Fig. 2.

**Figure 2:**
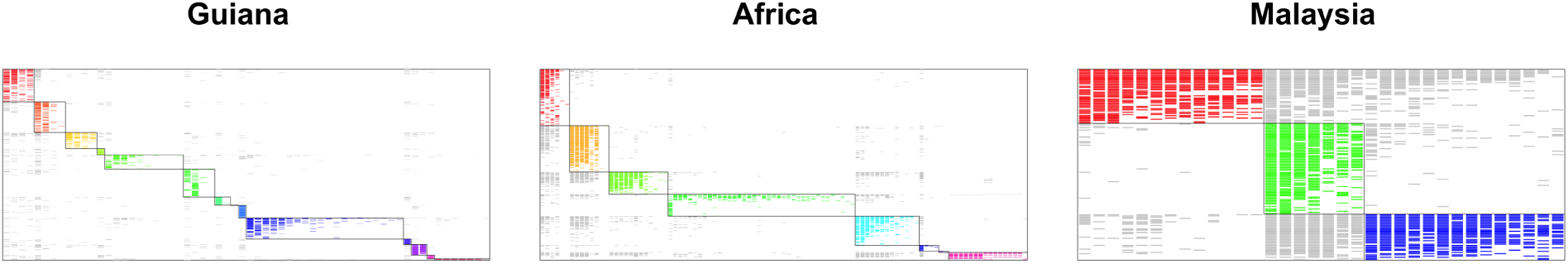
Adjacency matrices representing the three equatorial frugivory networks. Frugivore species are shown in the columns, plant species in the rows. Cell colors represent the modules identified in each network, which are also highlighted by contours. Grey cells represent interactions that do not belong to any particular module.

That was also the region with the highest modularity (Q = 0.48, 14 modules), significantly higher than expected under the free null model (mean = 0.37, P *<* 0.001). This indicates that its interactions are structured into distinct subsets of frugivore–plant associations. The nestedness at the entire matrix level (NODF = 14.33) was lower than expected by the free null model (mean = 14.74, P *<* 0.001), though slightly higher than expected by the restricted null model (mean = 12.73, P *<* 0.001). The nestedness within modules (NODF*_sm_* = 40.69) was higher than between modules (NODF*_dm_* = 11.00), in the former case higher than expected under the restricted null model (mean = 12.32, P *<* 0.001) and in the latter lower than expected (mean = 15.04, P *<* 0.001). Together, these results indicate a network dominated by a modular structure, with internally nested modules and limited cross-module overlap. In other words, they point to a compound topology; see Fig. 3. Centrality metrics revealed a small but distinct subset of frugivores that act as connectors or hubs, linking otherwise independent modules. Among frugivores, *Ramphastos tucanus* and *Ateles paniscus* emerged as connectors, bridging interactions across plant assemblages, while *Alouatta seniculus* and *Cebus apella*, for instance, acted as module hubs with high degree and eigenvector centralities. Plant species displayed a higher centrality than frugivore species, several of which were identified as module hubs or network hubs. *Ficus insipida* and *Symphonia globulifera*, for example, were identified as connectors, each linking multiple frugivore guilds. The relationship between the body mass of the frugivore and centrality was positive and significant in all metrics (*ρ* = 0.45–0.56, P*<*0.001), indicating that larger species tend to occupy more central positions. This pattern suggests that large frugivores contribute disproportionately to maintaining cross-module connectivity and potentially enhance long-distance seed dispersal within the Guianan network, Fig. 4.

**Figure 3:**
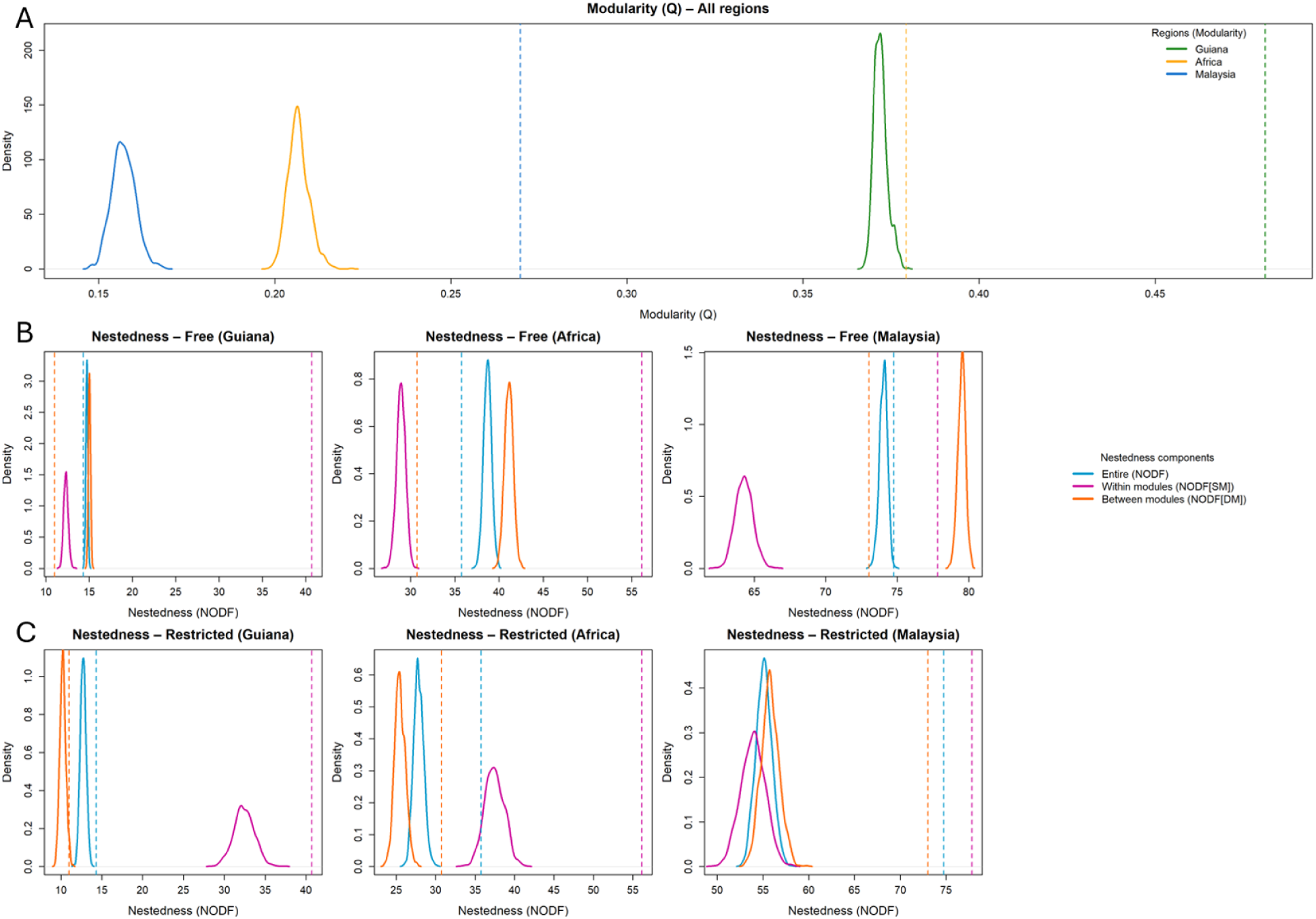
Distribution of randomized values for Modularity (*Q*) (panel A), Nestedness (NODF), within-module Nestedness (*NODF_SM_*), and between-module Nestedness (*NODF_DM_*) under the Free Null Model (panel B) and the Restricted Null Model (panel C), generated using Monte Carlo simulations for the Guianas, Africa and Malaysia networks. Dashed lines indicate the observed values of each metric.

**Figure 4:**
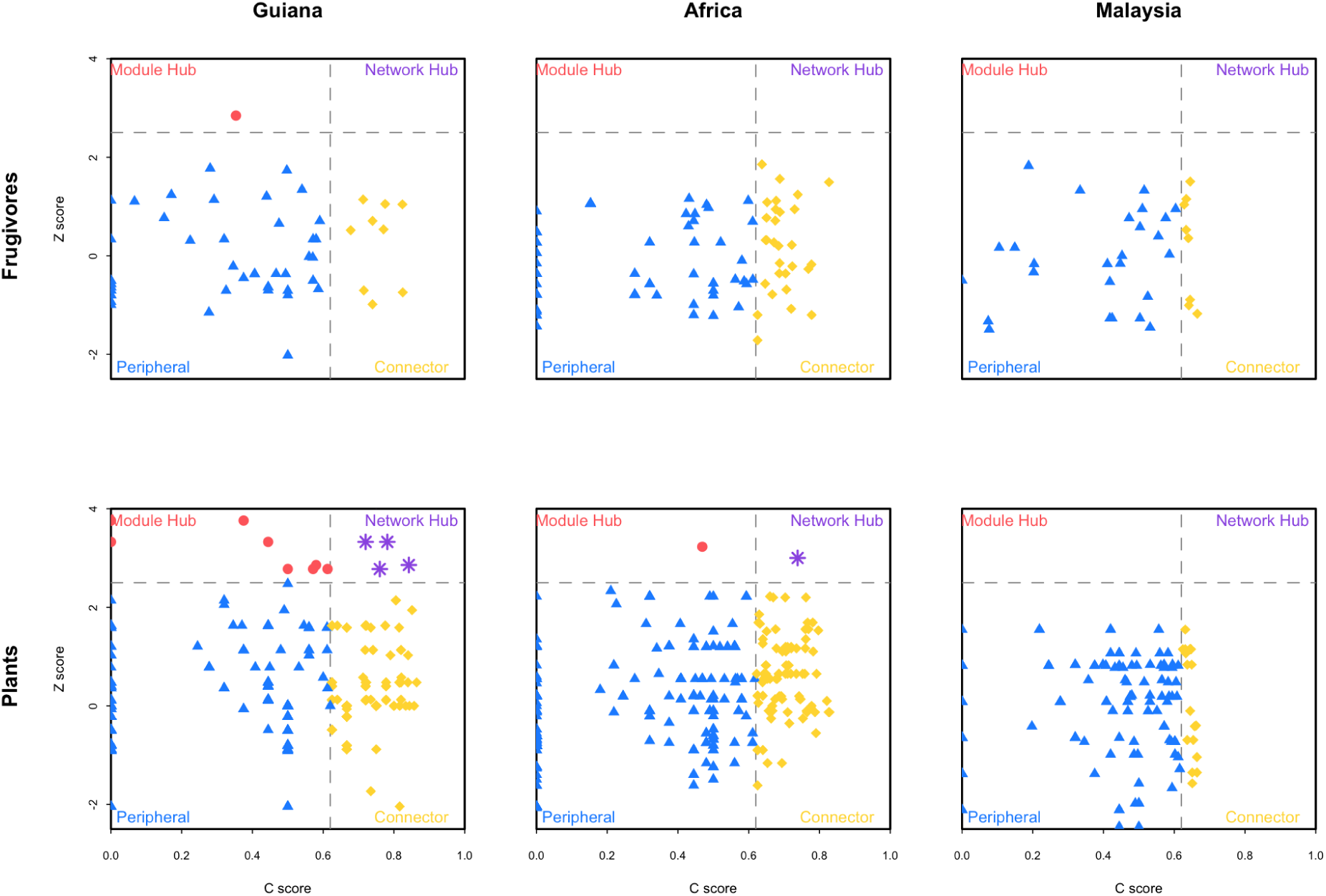
Functional roles based on *c* and *z* scores, detected for the frugivore and plant species of each equatorial network.

The composition of the Guianan modules is diverse, as is the composition of the frugivores and plant communities. Some modules contained both birds and mammals, whereas others were dominated by mammals. Bats were concentrated mainly in a nearly isolated module associated with Piperaceae and Moraceae, especially small-seeded *Piper* and *Ficus* species. Modules containing primates were mainly associated with Fabaceae, Sapotaceae, Moraceae, and Urticaceae. Terrestrial and understorey birds were associated with plants from understorey and open habitats whose fruits often fall to the ground, while manakins and dung beetles were associated with small-fruited and small-seeded plants, including *Psychotria*. These patterns indicate that the high modularity of the Guianan network reflects not only taxonomic turnover, but also differences in fruit size, seed size, handling behavior, and foraging strata.

### 3.2 Africa

The Central Africa network exhibited intermediate diversity, with 399 plant species, 99 frugivore species, and a total of 498 taxa. The turnover of species with the other regions was high (*β_plant_* = 0.99 vs. Guianas; *β_plant_* = 1.00 vs. Malaysia; *β_frugivore_* = 1.00 vs. Guianas; *β_frugivore_* = 0.97 vs. Malaysia), indicating strong regional distinctiveness in species composition.

Modularity was intermediate (Q = 0.379, 8 modules) and significantly higher than expected by chance (mean = 0.21, P *<* 0.001). The nestedness at the matrix level was moderate (NODF = 35.76), significantly lower than expected in the free null model (mean = 38.70, P*<*0.001) and higher than expected in the restricted model (mean = 27.85, P *<* 0.001). The nestedness was higher within (NODF*_sm_* = 56.14) than between modules (NODF*_dm_* = 30.74), and in both cases it was higher than expected under the restricted null model (means = 37.45 and 25.45, P*<*0.001), indicating that the African network is structured into modular clusters that are themselves internally nested. Such an organization suggests a balance between local compartmentalization and hierarchical integration within modules. In other words, these results point to a compound topology.

The centrality was balanced between plants and frugivores. Megafauna and other large-bodied frugivores, including *Loxodonta cyclotis*, *Mandrillus sphinx*, and *Colobus satanas*, were identified as connectors that link otherwise distinct plant modules. Several avian species, such as *Bycanistes albotibialis* and *Ceratogymna atrata*, functioned as hubs, concentrating many within-module interactions. On the plant side, *Symphonia globulifera* and *Myrianthus arboreus* appeared as key connectors shared with the Guiana network, indicating a conserved functional role across continents. The effect of body mass on centrality was consistently positive and highly significant in all metrics (*ρ* = 0.40–0.56, P*<*0.001), confirming that large-bodied frugivores tend to have higher degree, betweenness, and eigenvector centralities. This supports the idea that megafrugivores and large primates serve as keystone connectors that enhance network cohesion and resilience in the African assemblage.

The Africa modules also revealed a marked influence of natural-history. Some modules were dominated by primates, including great apes and large monkeys, and were mainly associated with Annonaceae, Fabaceae, Ebenaceae, Sapotaceae and Anacardiaceae. A module composed mainly of rodents was associated with Myristicaceae and likely represents the secondary handling of seed from seeds that fall to the ground after fruit consumption by other frugivores. Bird-dominated modules were more clearly separated from mammal modules than in the Guianas, and some of their dominant plant families, such as Apocynaceae and Euphorbiaceae, were not dominant in other modules. This indicates that the African network combines megafauna-driven connectivity with a clearer taxonomic and functional separation between avian and mammalian frugivory.

### 3.3 Malaysia

The Malaysia network was the least species-rich among the three regions, comprising 162 plant species, 34 frugivore species, and a total of 196 taxa. This richness was markedly lower than in Central Africa (498 species) and especially the Guianas (722 species). The species turnover was complete for plants compared to both Central Africa (*β_plant_* = 1.00) and the Guianas (*β_plant_* = 1.00) and almost complete for frugivores (*β_frugivore_* = 0.97 vs. Central Africa; *β_frugivore_* = 1.00 vs Guiana). These values confirm the strong biogeographic isolation of the Southeast Asian assemblage, which remains compositionally unique despite its lower overall richness within the pantropical gradient of frugivory networks.

Modularity was low but significant (Q = 0.27, 3 modules), higher than expected in the free null model (mean = 0.16, P*<*0.001). In contrast to the Guianas, nestedness was the dominant structural feature. The nestedness at the matrix level was high (NODF = 74.75) and exceeded the null expectations in both free and restricted models (means = 74.04 and 55.11, P*<*0.001). The nestedness was similar both within (NODF*_sm_* = 77.83) and between modules (NODF*_dm_* = 73.02), although it was lower than expected between modules (P*<*0.001) and higher than expected within modules (P *<* 0.001). These results describe a highly nested network in which highly connected species, both plants and frugivores, subsume the interaction sets of least-connected taxa, producing a hierarchically integrated architecture.

Centrality patterns were dominated by plant hubs rather than frugivores. Species such as *Ficus benjamina*, *Ficus racemosa*, and *Syzygium cumini* exhibited exceptionally high degree and betweenness centralities, confirming their roles as multi-frugivore hubs that anchor the nested network architecture. Among frugivores, the flying foxes *Pteropus vampyrus* and *Cynopterus brachyotis* were the main connectors, interacting with plant clusters across all modules. No significant correlation was found between the body mass of the frugivore and any centrality metric (*ρ <* 0.1, P *>* 0.3), reflecting a system where functional prominence is not tied to size, but rather to behavioral flexibility and resource breadth. In general, Malaysian network cohesion is sustained by structural hubs of plants and a few medium-sized generalist frugivores that maintain connectivity across the highly nested web.

The Malaysia network was organized into three modules, with birds separated from mammals. One module contained only birds, whereas the other two contained mammals. Moraceae were important across the network, but the dominant fruit types differed between modules: bird interactions were mainly associated with small-fruited *Ficus*, whereas mammal interactions were more strongly associated with large-fruited taxa such as *Artocarpus*. Euphorbiaceae were especially important in the bird module, whereas Annonaceae and Meliaceae were more important in the mammal modules. This organization suggests niche separation between birds and mammals, but also strong overlap through shared plant resources, consistent with the high nestedness observed in this network. Because several Malaysian frugivores are represented as groups rather than species, these patterns should be interpreted as reflecting both ecological organization and the taxonomic resolution of the dataset.

## 4 Discussion

Our pantropical comparison revealed structural and functional differences between the studied frugivory networks. Nevertheless, despite significant differences in species richness and composition, all networks shared a common qualitative pattern: a compound topology consistent with the Integrative Hypothesis of Specialization (IHS, (Mello and Dormann, 2025)). The balance between modularity and nestedness, as well as the identity of central species, varied across continents, reflecting their unique evolutionary and ecological histories. The Guianas and Africa networks were highly modular with internally nested modules, while the Malaysia network has a less marked archetypal structure, which still resembles a compound topology. Similarly, some species of large-bodied frugivores and plants served as connectors in the Guianas and Central Africa, while mainly plants played that role in Malaysia. These results suggest that the topology of frugivory networks is shaped by biogeographic history, body-size distributions, and the persistence or loss of megafauna, supporting the IHS view that different processes operate at different organizational scales, with resource heterogeneity as their main driver.

This gradient aligns with broader patterns of angiosperm and vertebrate radiation, with tropical South America retaining the greatest taxonomic and functional diversity (Carlucci et al., 2017). However, a higher species richness in the Guianas did not lead to greater interaction overlap, but rather to stronger partitioning between guilds, indicating that diversification favored specialization over redundancy (Dugger et al., 2019). In contrast, the Malaysia network, though species-poor, was anchored by a few generalist taxa that integrated the system (Ong et al., 2022). These differences show how evolutionary history and body-size spectra influence not only frugivore diversity but also the structure of their interactions with plants. They also show that network size matters, as larger systems are more likely to fit one of the four archetypal topologies, especially the compound topology.

Comparing network topologies across continents reinforces this view. In Guiana, where megafauna disappeared in the late Pleistocene (Galetti et al., 2017), the frugivory network showed strong modularity with nestedness within modules, forming a marked compound topology. Each module corresponded to a functional guild centered on a frugivore group (primates, toucans, or bats), suggesting that niche differentiation and resource heterogeneity were key structural drivers (Mello et al., 2019). The expected compensatory role of rodents through synzoochory (Ong et al., 2022) was partially supported, but not in a way that fully replaces the structural role of extinct megafrugivores. Dasyproctids, such as agoutis and acouchis, can act as module connectors because they handle both large and small seeds collected from the ground, providing secondary seed movement after fruits have fallen or after other frugivores have consumed the pulp. However, this role differs from that of megafauna because it is based on fruit handling and seed caching rather than the ingestion of the entire of part of large fruit and long-distance transport of large seeds. Thus, small mammals may contribute to local and secondary seed movement without necessarily restoring the cross-module connectivity formerly provided by extinct megafrugivores.

In Africa, where elephants and great apes still disperse seeds (Durand-Bessart et al., 2023), the network showed intermediate modularity and nestedness, reflecting a balance between local compartmentalization and regional integration (Thébault and Fontaine, 2010). Large frugivores bridged modules that might otherwise remain isolated (Sivault et al., 2023). In Malaysia, the network was predominantly nested, with a few highly connected species, especially figs and flying foxes, that linked most partners (Ong et al., 2022). Despite its lower modularity, the pattern of higher nestedness within than between modules indicates a weak compound topology dominated by an overarching nested core. This result diverges from our original prediction that the persistence of megafauna would drive cohesion in Southeast Asia. Instead, co-occurring *Ficus* and *Syzygium* species, coupled with the behavioral flexibility of flying foxes, appear to sustain network integration. This points to a distinct mechanism of cohesion, where resource phenology and plant redundancy replace the structural role of large-bodied dispersers. However, because the present analysis reflects a frugivory network rather than realized seed dispersal, the functional importance of animal dispersers may be underestimated. Frugivory networks capture fruit consumption patterns, while effective seed dispersal also depends on processes such as seed handling, intestinal passage, and deposition in suitable microsites (Beckman et al., 2023). Consequently, future network analyzes that explicitly incorporate seed dispersal effectiveness (Schupp et al., 2010) may reveal a stronger structural contribution of large-bodied frugivores than those observed here.

The natural-history composition of the modules helps explain why the same broad topological pattern can arise from different ecological mechanisms. In the Guianas, the near isolation of the bat module, associated with Piperaceae and small-seeded Moraceae, contrasts with primate modules associated with Fabaceae, Sapotaceae, Moraceae and Urticaceae. In Africa, primate and megafauna modules share functionally similar plant families, including Annonaceae, Fabaceae, Ebenaceae, Sapotaceae and Anacardiaceae, while bird modules are more distinct and associated with plant families such as Apocynaceae and Euphorbiaceae. In Malaysia, birds and mammals were more clearly separated into different modules, with birds associated primarily with small-fruited *Ficus* and Euphorbiaceae, while mammals were associated with Annonaceae, Meliaceae, and large-fruited Moraceae such as *Artocarpus*. These examples show that modularity reflects resource dissimilarity translated as differences in fruit traits, seed size, foraging strata, taxonomic identity, and handling behavior, rather than geography alone.

The contribution of megafauna to modularity and overall topology was not uniform between large-bodied species. Elephants acted as connector modules in Africa and Southeast Asia, and gorillas, chimpanzees, and mandrills also played connector roles in Africa. Nevertheless, some large Southeast Asian frugivores, such as gaur, Malayan tapir, and sun bear, were peripheral in the network. This may be related to fruit and seed handling: despite their size, these animals can chew or destroy seeds rather than swallow them whole (Ong et al., 2022). Therefore, body size alone is not sufficient to predict the role of the network. The ability to ingest, handle, transport, or cache seeds, together with resource breadth and taxonomic resolution of the data, seems to determine how a frugivore contributes to network cohesion. In Guiana and Africa, large-bodied frugivores such as howler monkeys, atelines, elephants, and mandrills occupied the most central positions, confirming that body size correlates with centrality (Sivault et al., 2023). Their ability to transport large seeds over long distances and interact with diverse plant assemblages makes them keystones that maintain network cohesion (Dennis et al., 2007). However, in Malaysia, centrality was highest in plants. Species such as *Ficus benjamina*, *F. racemosa*, and *Syzygium cumini* acted as structural hubs, while medium-sized bats served as secondary connectors (Albert-Daviaud et al., 2022). The lack of correlation between body size and centrality in this system suggests that behavioral flexibility and resource breadth, rather than morphology, determine prominence. Thus, in Southeast Asia, the stability of the frugivory network seems to depend more on abundance than consumer mobility, reversing the pattern seen in the Neotropics and Africa.

A second important limitation is the organization of the Southeast Asian database. Unlike the Guianan and African networks, in which frugivores are identified mainly at the species level, several animals in the Southeast Asian network represent groups of closely related species, including rats, squirrels, hornbills, bats, and bulbuls (Ong et al., 2022). This may overestimate the apparent role of these grouped taxa and, at the same time, underestimate the network modularity and specialization of the network. A finer taxonomic resolution could increase the number of modules and reduce nestedness, possibly making the Southeast Asian network more similar to the African network. Finally, the differences between our results and those of previous analysis of the Malaysian network may also result from methodological choices. Previous work used quantified data and Newman’s modularity, whereas the present comparison used binary matrices and the same bipartite modularity algorithm for all networks. This standardization improves comparability between regions, but it necessarily simplifies local ecological detail.

The present study should be interpreted as a structural comparison of frugivory networks, not as a direct comparison of the realization of seed dispersal. Effective dispersal depends on additional processes that were not available for all regions, including seed survival after fruit handling, intestinal passage, deposition site, post-dispersal predation, and seedling recruitment Schupp et al. (2010). The patterns reported here therefore describe the organization of frugivory interactions and identify plausible mechanisms by which biogeographic history, megafauna, plant traits, and frugivore behavior may ultimately influence seed dispersal and forest dynamics.

Together, these findings offer a biogeographic test of IHS applied to frugivory and seed dispersal (like previusly done for resin collection, Nakamura et al., 2024). Across continents, frugivory networks tend toward a compound topology, but the relative influence of modular and nested components shifts with biogeography, evolution, and ecology. IHS proposes that large-scale phylogenetic and biogeographic constraints shape the modularity of ecological networks, while resource breadth drives nestedness within modules (Mello et al., 2019). Our results support these predictions. In the Guianas, diversification and ecological segregation produced a typically compound topology. In Africa, megafrugivores integrated different guilds into a less modular network. In Malaysia, a few generalist plants and bats dominated a more nested structure shaped by resource overlap. At the same time, the comparison also shows that small and medium-sized frugivores can have important and complementary roles when they are able to handle large fruits and seeds and connect modules through secondary seed movement. However, because available datasets remain uneven across regions and taxa, especially for rodents, bats, and poorly identified frugivore groups, these conclusions should be treated as new hypotheses to be refined with more balanced sampling and higher taxonomic resolution. This pantropical gradient shows that the tropical frugivory architecture is not random but emerges from the layered influence of multiple processes that act on multiple scales.

## 5 Supplementary Material

Detailed Supplementary Methods are available with the online version of this article. The complete annotated source code and operational workflow documentation are available in the associated GitHub repository. The repository provides methodological transparency and supports computational reproducibility for users with authorized access to the original data; it does not contain the source data, intermediate analytical results, or generated figures.

